# Epidermal CD109 Overexpression Limits Cutaneous Inflammatory Signaling

**DOI:** 10.64898/2026.03.13.711666

**Authors:** Adel Batal, Jean-Philip Lacroix, Joshua Vorstenbosch, Melani Lighter, Anie Philip

## Abstract

Psoriasis is a chronic immune-mediated inflammatory skin disease characterized by excessive keratinocyte proliferation, immune cell infiltration and dysregulated inflammatory signaling. Despite the availability of biologic therapies targeting inflammatory cytokines, many patients experience incomplete responses or relapse, highlighting the need to better understand molecular regulators of cutaneous inflammation. CD109 is a glycosylphosphatidylinositol (GPI)-anchored protein previously identified by our lab as a co-receptor and negative regulator of Transforming Growth Factor-β (TGF-β) signaling that inhibits fibrotic responses. Emerging evidence suggests that CD109 also modulates immune and inflammatory pathways. In this study, we investigated whether epidermal CD109 overexpression influences cutaneous inflammatory responses. Transgenic (TG) mice overexpressing CD109 under the keratin-14 (K14) promoter were used to restrict transgene expression to the epidermis. TG and wild-type (WT) littermates were subjected to lipopolysaccharide (LPS)-induced skin inflammation. CD109 TG mice exhibited significantly reduced immune cell recruitment, including macrophages and neutrophils, along with decreased expression of the pro-inflammatory mediators IL-1α and MCP-1/CCL2 compared with WT mice. Transcriptomic analysis of primary keratinocytes revealed downregulation of multiple inflammatory signaling pathways in CD109-overexpressing cells, including TNF-α/NF-κB, IL-2/STAT5, IFN-γ, IFN-α, and IL-6/JAK/STAT3 pathways. Together, these findings demonstrate that epidermal CD109 overexpression attenuates cutaneous inflammatory responses by suppressing key inflammatory signaling networks and limiting immune cell recruitment, suggesting that CD109 may represent an important regulator of inflammatory signaling in the skin and a potential target for inflammatory skin diseases such as psoriasis.

## INTRODUCTION

Psoriasis is a chronic immune-mediated inflammatory skin disease that affects approximately 2-3% of the global population, making it one of the most common inflammatory skin disorders worldwide [1]. The disease is characterized by excessive keratinocyte proliferation, epidermal hyperplasia, and infiltration of immune cells, leading to erythematous, scaly plaques that can significantly impair quality of life [2]. Beyond the skin, psoriasis is associated with several systemic comorbidities, including cardiovascular disease, metabolic syndrome, and psoriatic arthritis, which collectively contribute to increased morbidity and mortality among affected individuals [2, 3]. Despite the availability of topical therapies, systemic immunosuppressants, and biological agents targeting inflammatory cytokines such as TNF-α, IL-17, and IL-23, many patients experience incomplete responses, adverse effects, or relapse after treatment discontinuation [2, 3]. These limitations highlight the need to better understand the molecular mechanisms that regulate inflammatory signaling in the skin to identify new therapeutic targets.

CD109 is a glycosylphosphatidylinositol (GPI)-anchored protein that our lab has identified as a Transforming Growth Factor Beta (TGF-β) co-receptor that inhibits TGF-β signaling and fibrotic responses [4-9]. CD109 is increasingly recognized as a crucial regulator of immune and inflammatory responses in various tissues and cell types [10].

Growing evidence suggests that CD109 plays an important role in regulating inflammatory signaling networks. Batal et al. described CD109 as a key regulator of inflammatory responses across multiple tissues, including the skin, lung, bone, and immune cells [10]. In addition to modulating TGF-β signaling, CD109 appears to influence several pathways that are central to the control of inflammatory responses [10]. These include Nuclear Factor Kappa B (NF-κB) signaling, interferon-mediated responses, and cytokine-driven pathways, such as JAK/STAT signaling, which collectively regulate the expression of pro-inflammatory mediators including cytokines, chemokines, and other immune regulators [10].

In the skin, keratinocytes play a central role in orchestrating inflammatory responses. Beyond serving as a physical barrier, keratinocytes actively participate in immune regulation by producing cytokines and chemokines that recruit and activate immune cells within the dermal compartment [11]. Through these signaling interactions, keratinocytes help coordinate the initiation and amplification of cutaneous inflammatory responses following tissue injury or microbial challenge [11, 12]. Consequently, molecules that regulate inflammatory signaling within keratinocytes may profoundly influence the magnitude and progression of inflammation in the skin. Based on the emerging role of CD109 as a regulator of inflammatory signaling pathways, we hypothesize that CD109 overexpression may suppress cutaneous inflammation by modulating key inflammatory signaling networks in keratinocytes.

## METHODS

### LPS-Induced Inflammation Model

Transgenic mice (TG) overexpressing CD109 under the control of the K14 promoter to restrict transgene expression to the epidermis have been generated [13-15]. CD109 TG mice and their wild-type (WT) littermates (aged 6-8 weeks) were anesthetized by isoflurane (Sigma-Aldrich, Mississauga, ON), shaved on their dorsal surface and depilated using Nair (Church & Dwight,York, PA). Mice were injected intradermally with 5 μg of filter-sterilized lipopolysaccharide (LPS, Sigma Aldrich, Mississauga, ON) in phosphate buffered saline (PBS) or with PBS alone, into a single (1 cm2) site on their shaved backs. Mice were sacrificed by CO2 asphyxiation at 24 or 48 hours post-injection and the injected skin tissue was harvested, bisected, and either fixed in 10% neutral buffered formalin (Sigma-Aldrich) for histological/immunohistochemical analysis or snap-frozen in liquid nitrogen and stored at -80°C (for total RNA extraction and qPCR analysis). All animal experiments were approved by the McGill University Animal Care Committee.

### Histology and Immunohistochemistry

Formalin-fixed skin tissue was embedded in paraffin and cut into 7-μm sections using a microtome. Sections were stained with hematoxylin and eosin (H & E) for microscopic evaluation using ImageProPlus6 Software (Media Cybernetics). For immunohistochemistry, antigen retrieval was performed by boiling slides in sodium citrate buffer (10 mM sodium citrate, 0.05% Tween 20, pH 6.0) for 10 minutes. Slides were washed and blocked using 5% normal goat serum (NGS) and then incubated overnight at 4°C with anti-F4/80 (macrophage marker) or Ly-6G (Gr-1, neutrophil marker) antibodies (eBioscience, San Diego, CA) diluted 1:50 in 5% NGS. The following day, slides were washed and incubated with a biotinylated secondary antibody, followed by ABC reagent (Vector Labs, Burlington, ON, Canada) and then developed using ImmPACT DAB (Vector Labs). The number of macrophages and neutrophils was quantified by taking the average number of F4/80 or Ly-6G positive cells per high power field.

### Quantitative Real-Time PCR

Total RNA was extracted from skin by homogenization in Trizol (Invitrogen) and purified using an RNEasy Kit following the manufacturer’s protocol (Qiagen). Reverse transcription was performed using MMLV Reverse Transcriptase (Invitrogen). 1 μg of RNA was reverse transcribed using oligo-dT primer in a final volume of 20 μL. qPCR was performed to amplify interleukin-1α (IL-1α) (Forward Primer: 5′-TCGGGAGGAGACGACTCTAA-3′; Reverse Primer: 5′-GTATCATATGTCGGGGTGGC-3), monocyte chemoattractant protein-1 (MCP-1) (Forward Primer: 5′-CATGCTTCTGGGCCTGCTGTTC-3′; Reverse Primer: 5′ CATGCTTCTGGGCCTGCTGTTC-3′) and glyceraldehyde 3-phosphate dehydrogenase (GAPDH) housekeeping control (Forward Primer: 5′-GGCGTCTTCACCACCATGGAG-3′; Reverse Primer: 5′-AAGTTGTCATGGATGACCTTGGC-3′) with iQ SYBR Green Supermix on a CFX96 Thermocycler (Bio-Rad, Hercules, CA). Relative mRNA levels were calculated using the ΔΔCt method according to the formula ΔΔCt = ΔCt (GOI_treatment_ - GAPDH_treatment_) - ΔCt (GOI_control_ - GAPDH_control_). GOI = gene of interest, and the data are presented as fold-change (2ΔΔCt) [16].

### Keratinocyte Isolation and Culture

Primary mouse keratinocytes were isolated from the skin of newborn CD109 TG mice and WT littermates. Briefly, full-layer skin was removed from newborn (1–2 days old) mice and treated with 0.25% trypsin (Invitrogen) overnight at 4°C. The epidermis was then separated from the dermis, minced into smaller pieces, placed in a sterile 15-ml tube containing 5 ml of low calcium EMEM media and agitated to create a single-cell suspension. The cells were resuspended in fresh media and plated in culture dishes coated with type IV collagen for attachment. Cells were incubated at 37°C under 5% CO2 in a humidified incubator and media were changed every 2-3 days.

### Microarray Analysis

Total RNA was extracted from primary CD109 transgenic (n=3) and wild-type (n=3) keratinocytes using the RNeasy Mini Kit (Qiagen, Mississauga, ON) and RNA integrity was verified using a Bioanalyzer (Agilent). Total RNA (250 ng) was amplified using the TotalPrep RNA Amplification kit (Ambion) and the biotinylated cRNAs were hybridized to the MouseWG-6 v2.0 Expression BeadChip (Illumina Inc., San Diego, CA). The multi-sample format allowed us to profile more than 45,200 transcripts for 6 different samples (3 TG versus 3 WT) simultaneously on a single BeadChip. Hybridized biotinylated cRNA was detected using Cy3-conjuated streptavidin and fluorescence emission by Cy3 was quantified using an iScan reader (Illumina Inc.). Data were analyzed using FlexArray Software V1.6.1 (McGill University and Genome Quebec Innovation Center, Montreal, QC, http://genomequebec.mcgill.ca/FlexArray/license.php). The Microarray data were subsequently used for gene set enrichment analysis (GSEA) to visualize and compare the enrichment of various gene sets between TG and WT samples.

## RESULTS

### CD109 overexpression in the epidermis inhibits immune cell recruitment during lipopolysaccharide (LPS)-induced inflammation in the skin

CD109 TG mice show reduced macrophage (p<0.05) and neutrophil (p<0.01, p<0.001) recruitment compared to WT littermates in response to LPS-induced skin inflammation **(Figure 1A, 1B)**. Quantification of F4/80-positive macrophages and Ly6G-positive neutrophils demonstrated a significant reduction in immune cell infiltration in TG skin at both 24 and 48 hours following LPS injection. This was evidenced by decreased dermal cellularity in H&E-stained sections **(Figure 1C)** and lower immune cell counts in TG mice at 24 and 48 hours post-injection.

**Figure 1:**
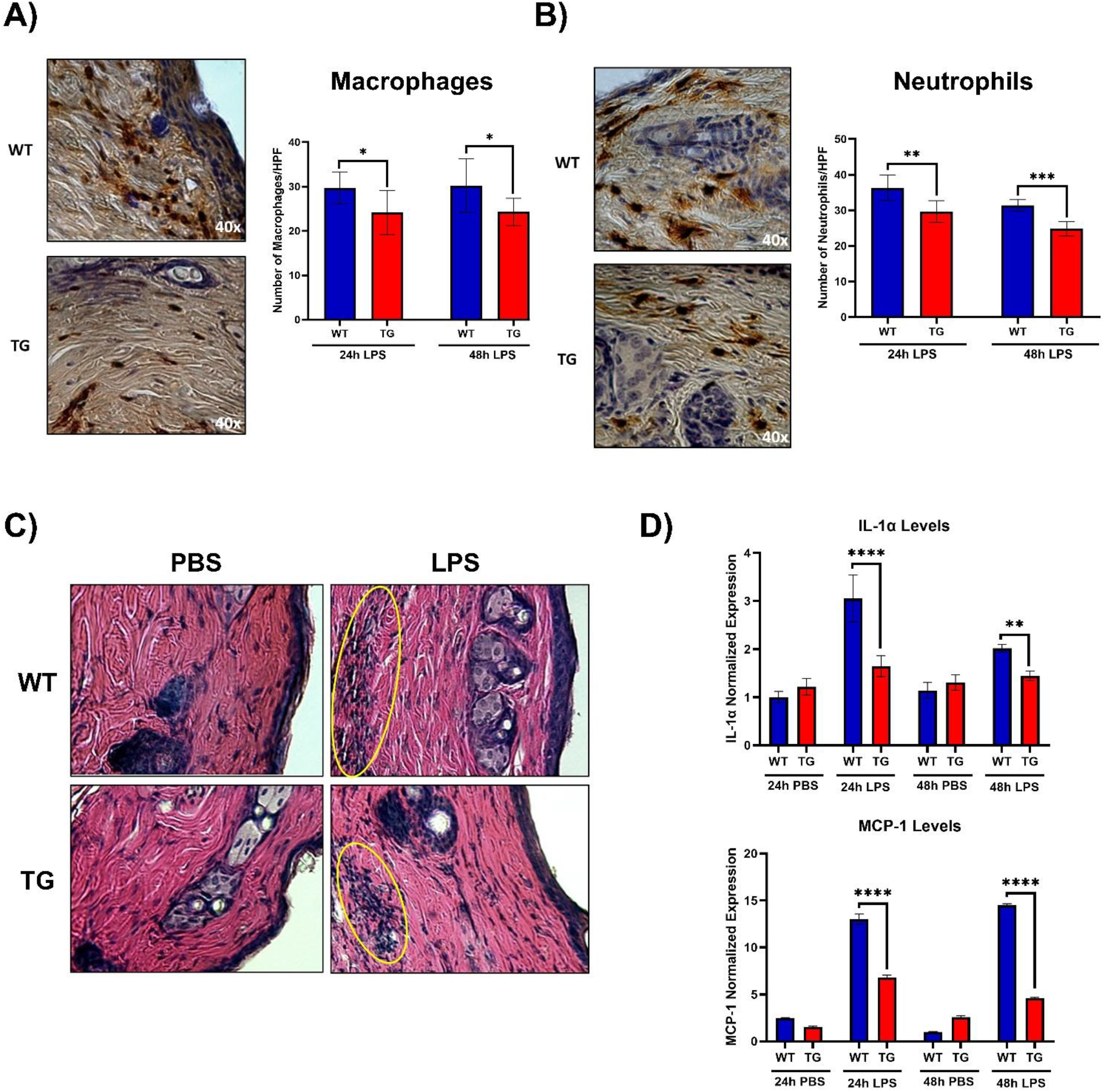
Epidermal overexpression of CD109 reduces immune cell infiltration, and inflammatory cytokine expression. **A)** Quantification of macrophage (F4/80) and **B)** neutrophil (Ly6G) infiltration in wild-type (WT) and CD109 transgenic (TG) mouse skin at 24 and 48 hours after intradermal LPS injection. Representative immunohistochemical staining images (left panels) show reduced immune cell presence in TG skin. Bar graphs (right panels) show mean ± SEM of positive cells per mm^2^ dermis (n = 6 per group). *p < 0.05, **p < 0.01, ***p < 0.001. **C)** Representative H&E-stained skin sections from WT and TG mice 24 hours after injection with LPS or PBS control. TG skin displays decreased dermal cellularity compared to WT in response to LPS-induced inflammation. **D)** Relative mRNA expression of pro-inflammatory cytokines IL-1α and MCP-1/CCL2 in WT and TG skin at 24 and 48 hours post-LPS injection, measured by qPCR. Data are normalized to housekeeping genes and expressed as fold change relative to PBS controls. Bars represent mean ± SEM (n = 6 per group). *p < 0.05, **p < 0.01, ***p < 0.001.

### CD109 overexpression in the epidermis decreases pro-inflammatory cytokine (IL-1α and MCP-1/CCL2) mRNA expression during LPS-induced inflammation in the skin

CD109 TG mice exhibit a marked downregulation of pro-inflammatory cytokine expression in response to LPS-induced skin inflammation. IL-1α mRNA levels, which increase ∼3-fold in WT mice at 24 hours (p< 0.0001) and 48 hours (p<0.01) post-injection, show no significant change in TG mice **(Figure 1D upper panel)**. Likewise, MCP-1/CCL2 mRNA expression is significantly attenuated in TG mice, showing approximately a 3-fold decrease compared to WT levels at 24 hours (p<0.0001), followed by a further ∼50% reduction at 48 hours (p<0.0001) **(Figure 1D lower panel)**. These findings indicate that CD109 overexpression in keratinocytes suppresses the production of key cytokines and chemokines involved in immune cell recruitment during acute inflammation.

### CD109 overexpression in the epidermis downregulates pro-inflammatory and immune signaling pathways

Transcriptomic analysis of primary keratinocytes isolated from CD109 TG and WT mice revealed significant enrichment of multiple immune-related pathways. reveals significant enrichment of multiple immune-related pathways in TG mice compared to WT. These include hallmark inflammatory responses (FDR = 0.0031, **Figure 2A**), TNF-α signaling via NF-κB (FDR = 0.0017, **Figure 2B**), IL-2/STAT5 signaling (FDR = 0.0023, **Figure 2C**), IFN-γ signaling (FDR = 0.0017, **Figure 2D**), IFN-α signaling (FDR = 0.0023, **Figure 2E**), and IL-6/JAK/STAT3 signaling (FDR = 0.011, **Figure 2F**). Downregulated genes across these pathways include key inflammatory mediators and transcriptional regulators such as TLR2, TNF, IL1A, CCL2, NFKB1, IRF1, SOCS1, and STAT1, which are known to play critical roles in innate immune signaling and cytokine-mediated inflammatory responses.

**Figure 2:**
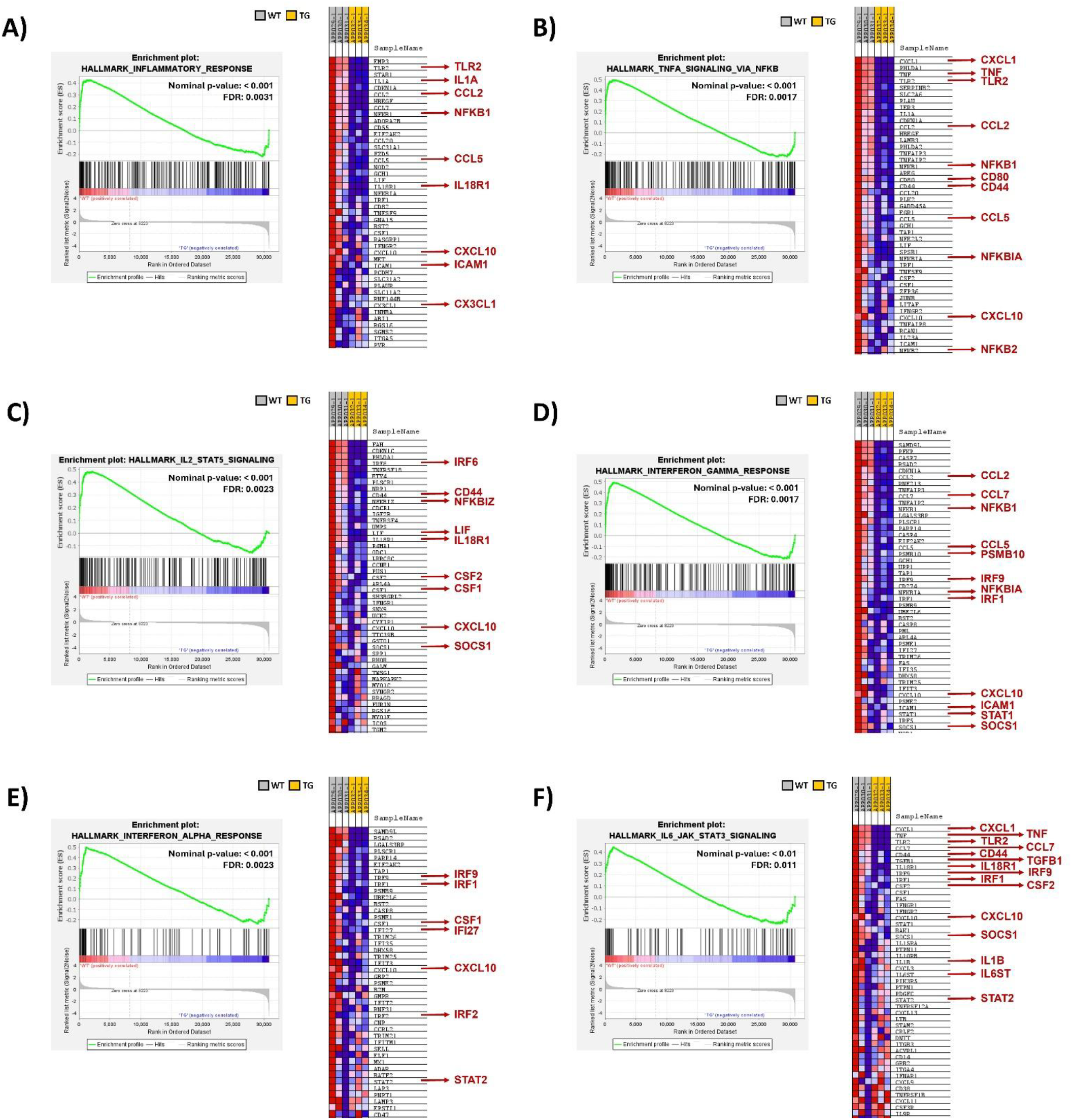
CD109 overexpression suppresses pro-inflammatory signaling pathways in keratinocytes. **A-F)** Gene Set Enrichment Analysis (GSEA) and associated heatmaps comparing gene expression profiles between WT and TG primary keratinocytes. Enrichment plots show negative enrichment of hallmark immune pathways in TG keratinocytes, including: A) Inflammatory Response (FDR = 0.0031), **B)** TNF-α signaling via NF-κB (FDR = 0.0017), **C)** IL-2/STAT5 signaling (FDR = 0.0023), **D)** IFN-γ response (FDR = 0.0017), **E)** IFN-α response (FDR = 0.0023), and **F)** IL-6/JAK/STAT3 signaling (FDR = 0.011). Heatmaps display expression levels of representative genes (red arrows) within each pathway, with yellow bars indicating TG samples and white bars indicating WT. Red = high expression; blue = low expression.

## DISCUSSION

Previous studies conducted by our team have shown that CD109 TG mice display reduced inflammation during cutaneous wound healing and bleomycin-induced skin fibrosis [13, 14], suggesting that CD109 displays anti-inflammatory properties *in vivo*. Here we show that CD109 TG mice display a decrease in immune cell (neutrophil and macrophage) numbers and diminished pro-inflammatory cytokine (IL-1α and MCP-1/CCL2) mRNA levels *in vivo* as compared to WT littermates, in a cutaneous LPS-induced model of acute inflammation.

These findings are consistent with previous studies from our laboratory demonstrating that epidermal CD109 overexpression reduces inflammation and improves tissue repair during wound healing. In the study by Vorstenbosch et al. (2013), CD109 transgenic mice showed decreased inflammatory cell infiltration and improved collagen organization during cutaneous wound healing, highlighting the importance of CD109 in regulating inflammatory responses during tissue repair [13].

In addition, using global gene expression profiling, we have identified five downregulated inflammatory signaling pathways in the TG mice (TNF-α/NF-κB, IL-2/STAT5, IFN-γ, IFN-α, IL-6/JAK/STAT3).

Keratinocytes are known to regulate immune responses in the skin through the release of cytokines and chemokines that recruit immune cells to the dermis [11]. Therefore, the suppression of inflammatory gene expression observed in CD109-overexpressing keratinocytes likely contributes to reduced dermal immune cell infiltration through epidermal-dermal signaling interactions. This concept of epidermal-dermal crosstalk has been previously highlighted in studies showing that epidermal signaling molecules can profoundly influence inflammatory responses in the dermal compartment [17-19].

Our findings suggest that CD109 plays a multifaceted role in regulating immune and tissue repair processes by modulating both the TGF-β pathway and several key inflammatory signaling cascades. In addition to influencing NF-κB activity, CD109 appears to impact other cytokine-driven pathways. This broader regulatory profile points to a potential role for CD109 in orchestrating the crosstalk between pro-inflammatory responses and tissue regeneration mechanisms. These observations raise important questions about the mechanisms by which CD109 coordinates this balance, and how it affects immune cell dynamics, cytokine environments, and the resolution of inflammation during wound healing.

Together, these results support the idea that CD109 acts as an important regulator of cutaneous inflammatory signaling and may contribute to maintaining immune homeostasis in the skin by limiting excessive inflammatory responses.

The specific effects of CD109 within the dermis remain unclear and have yet to be thoroughly investigated. Further studies are needed to understand CD109’s potential anti-inflammatory role in the skin, notably in the dermis.

## REFERENCES

1. Ponikowska, M., et al., Challenges Psoriasis and Its Impact on Quality of Life: Challenges in Treatment and Management. Psoriasis (Auckl), 2025. 15: p. 175–183.

2. Gonçalves, M.B.S., et al., Advancing insights into psoriasis: from pathogenesis to current and emerging therapies. International Immunopharmacology, 2025. 165: p. 115429.

3. Kakarala, C.L., et al., Beyond the Skin Plaques: Psoriasis and Its Cardiovascular Comorbidities. Cureus, 2021. 13(11): p. e19679.

4. Man, X.Y., et al., CD109, a TGF-β co-receptor, attenuates extracellular matrix production in scleroderma skin fibroblasts. Arthritis Res Ther, 2012. 14(3): p. R144.

5. Finnson, K.W., et al., Identification of CD109 as part of the TGF-beta receptor system in human keratinocytes. Faseb j, 2006. 20(9): p. 1525–7.

6. Bizet, A.A., et al., The TGF-β co-receptor, CD109, promotes internalization and degradation of TGF-β receptors. Biochim Biophys Acta, 2011. 1813(5): p. 742–53.

7. Bizet, A.A., et al., CD109-mediated degradation of TGF-β receptors and inhibition of TGF-β responses involve regulation of SMAD7 and Smurf2 localization and function. J Cell Biochem, 2012. 113(1): p. 238–46.

8. Li, C., et al., Soluble CD109 binds TGF-β and antagonizes TGF-β signalling and responses. Biochem J, 2016. 473(5): p. 537–47.

9. Litvinov, I.V., et al., CD109 release from the cell surface in human keratinocytes regulates TGF-β receptor expression, TGF-β signalling and STAT3 activation: relevance to psoriasis. Exp Dermatol, 2011. 20(8): p. 627–32.

10. Batal, A., et al., CD109, a master regulator of inflammatory responses. Front Immunol, 2024. 15: p. 1505008.

11. Jiang, Y., et al., Cytokinocytes: the diverse contribution of keratinocytes to immune responses in skin. JCI Insight, 2020. 5(20).

12. Piipponen, M., D. Li, and N.X. Landén, The Immune Functions of Keratinocytes in Skin Wound Healing. Int J Mol Sci, 2020. 21(22).

13. Vorstenbosch, J., et al., Transgenic mice overexpressing CD109 in the epidermis display decreased inflammation and granulation tissue and improved collagen architecture during wound healing. Wound Repair Regen, 2013. 21(2): p. 235–46.

14. Vorstenbosch, J., et al., CD109 overexpression ameliorates skin fibrosis in mouse model of bleomycin-induced scleroderma. Arthritis Rheum, 2013. 65(5): p. 1378–83.

15. Vorstenbosch, J., et al., Overexpression of CD109 in the Epidermis Differentially Regulates ALK1 Versus ALK5 Signaling and Modulates Extracellular Matrix Synthesis in the Skin. Journal of Investigative Dermatology, 2017. 137(3): p. 641–649.

16. Schmittgen, T.D. and K.J. Livak, Analyzing real-time PCR data by the comparative C(T) method. Nat Protoc, 2008. 3(6): p. 1101–8.

17. Costello, L., et al., Investigation into the significant role of dermal-epidermal interactions in skin ageing utilising a bioengineered skin construct. J Cell Physiol, 2025. 240(1): p. e31463.

18. Rebholz, B., et al., Crosstalk between keratinocytes and adaptive immune cells in an IkappaBalpha protein-mediated inflammatory disease of the skin. Immunity, 2007. 27(2): p. 296–307.

19. Ye, J. and Y. Lai, Keratinocytes: new perspectives in inflammatory skin diseases. Trends Mol Med, 2025. 31(12): p. 1103–1113.

